# Domain-specific cognitive impairment reflects prefrontal dysfunction in aged common marmosets

**DOI:** 10.1101/2023.05.22.541766

**Authors:** Courtney Glavis-Bloom, Casey R Vanderlip, Payton A Asch, John H Reynolds

## Abstract

Age-related cognitive impairment is not expressed uniformly across cognitive domains. Cognitive functions that rely on brain areas that undergo substantial neuroanatomical changes with age often show age-related impairment, while those that rely on brain areas with minimal age-related change typically do not. The common marmoset has grown in popularity as a model for neuroscience research, but robust cognitive phenotyping, particularly as a function of age and across multiple cognitive domains, is lacking. This presents a major limitation for the development and evaluation of the marmoset as a model of cognitive aging, and leaves open the question of whether they exhibit age-related cognitive impairment that is restricted to some cognitive domains, as in humans. In this study, we characterized stimulus-reward association learning and cognitive flexibility in young adults to geriatric marmosets using a Simple Discrimination and a Serial Reversal task, respectively. We found that aged marmosets show transient impairment in “learning-to-learn” but have conserved ability to form stimulus-reward associations. Furthermore, aged marmosets have impaired cognitive flexibility driven by susceptibility to proactive interference. Since these impairments are in domains critically dependent on the prefrontal cortex, our findings support prefrontal cortical dysfunction as a prominent feature of neurocognitive aging. This work positions the marmoset as a key model for understanding the neural underpinnings of cognitive aging.

**Significance Statement:** Aging is the greatest risk factor for neurodegenerative disease development, and understanding why is critical for the development of effective therapeutics. The common marmoset, a short-lived non-human primate with neuroanatomical similarity to humans, has gained traction for neuroscientific investigations. However, the lack of robust cognitive phenotyping, particularly as a function of age and across multiple cognitive domains limits their validity as a model for age-related cognitive impairment. We demonstrate that aging marmosets, like humans, have impairment that is specific to cognitive domains reliant on brain areas that undergo substantial neuroanatomical changes with age. This work validates the marmoset as a key model for understanding region-specific vulnerability to the aging process.

## INTRODUCTION

The effects of aging on cognitive function vary enormously from person to person. Some individuals retain high levels of cognitive ability throughout life, while others experience varying degrees of cognitive impairment (Nyberg et al., 2020). This cognitive impairment is not expressed uniformly across all cognitive domains. Cognitive functions that rely on brain areas that undergo substantial neuroanatomical changes with age often show age-related impairment, while those that rely on brain areas with minimal age-related change typically do not (Samu et al., 2017). To understand the processes that contribute to this variability in humans, it is critical to study model systems that share behavioral, neuroanatomical, and age-related neuropathological features with humans. As such, the common marmoset (*Callithrix jacchus*) has emerged as an advantageous model in which to investigate the biological consequences of aging, and they also have the advantage of having a short lifespan (∼10 years), which makes it feasible to undertake longitudinal studies examining the processes of aging over their entire lifespan (Tardif, 2019; Glavis-Bloom et al., 2023; Perez-Cruz and Rodriguez-Callejas, 2023).

Despite their growing popularity as a model, robust cognitive phenotyping of marmosets, particularly as a function of age and across multiple cognitive domains, is lacking. This presents a major limitation for the development and evaluation of the marmoset as a model of cognitive aging, and leaves open the question of whether they exhibit age-related cognitive impairment that is restricted to some cognitive domains, as in humans.

Two cognitive domains that, in humans, are differentially vulnerable to aging are stimulus-reward association learning and cognitive flexibility. Stimulus-reward association learning, which is dependent on the striatum, is typically assessed using simple discrimination tasks. On these tasks, through trial and error, animals learn which stimulus is associated with a reward. Existing literature shows that non-human primate performance on these tasks is unaffected by aging (Bartus et al., 1979; Burke et al., 2011), which aligns with the fact that the striatum undergoes only minimal changes with age (Cox et al., 2008; Samanez-Larkin et al., 2011). Cognitive flexibility, however, is a domain that is vulnerable to the effects of aging. Reversal learning tasks, which require animals to learn new stimulus-reward associations following uncued reinforcement shifts, are commonly used to quantify cognitive flexibility (Lai et al., 1995; Meunier et al., 1997). The critical involvement of the prefrontal cortex for cognitive flexibility is well established from lesion studies in marmosets, macaques, and humans (Clarke et al., 2004; Hornak et al., 2004; La Camera et al., 2018). Given that the prefrontal cortex undergoes significant changes with age (Resnick et al., 2007; Upright and Baxter, 2021; Glavis-Bloom et al., 2023), it is unsurprising that aged macaque monkeys and humans display significantly impaired cognitive flexibility compared to young controls (Bartus et al., 1979; Lai et al., 1995; Earles et al., 1997; Voytko, 1999; Head et al., 2009; Burke et al., 2011; Gray et al., 2023).

To date, a handful of studies have investigated stimulus-reward association learning and cognitive flexibility in aged marmosets, with mixed results. Two groups report that aged marmosets have intact stimulus-reward association learning, but impaired cognitive flexibility (Munger et al., 2017; Sadoun et al., 2019). However, another group found the opposite effects of aging, reporting that the ability to learn stimulus-reward associations declined with age, while cognitive flexibility did not (Rothwell et al., 2022). To reconcile these incongruent findings, we employed a serial reversal paradigm to increase proactive interference (Dickinson and Mackintosh, 1978; Hassett and Hampton, 2017). Using this approach, we probed the limits of cognitive flexibility as a function of age by escalating prefrontal cortical demand. Through detailed analyses of individual performance metrics, we find that aged marmosets have impaired cognitive flexibility that is explained by susceptibility to proactive interference.

## MATERIALS AND METHODS

### Subjects

Eleven (5 male, 6 female) common marmosets (*Callithrix jacchus*) between the ages of 3.4 and 10.9 years of age participated in this study. Animals younger than seven years of age were classified as “Young”, and those seven years of age or older were classified as “Aged”. Each of these marmosets was previously trained to perform cognitive tasks presented on touch screen computers installed in their home cages (Glavis-Bloom et al., 2022). In the present study, each of the 11 marmosets were tested on a Simple Discrimination task to measure their ability to make stimulus-reward associations. Seven of these animals (2 male, 5 female; 3.4-10.1 years old) were also tested on a Serial Reversal task to measure cognitive flexibility. Marmosets were tested for 1-3 hours on 3-5 days per week. They worked for fluid rewards, but were not food or water restricted.

Animals were either singly or pair housed in cages that contained enrichment items (e.g., hammocks and manzanita branches), and all animals had visual and auditory access to other marmosets. Home cages housing paired marmosets were split in half for the few hours during which cognitive testing was performed. All experimental procedures were carried out under a protocol that was approved by the Salk Institute Institutional Animal Care and Use Committee and conformed to NIH guidelines for the care and use of laboratory animals.

### Equipment

#### Touchscreen stations and cages

Marmoset home cages were custom-built to include a testing chamber in an upper corner of the cage, onto which a 10.4-inch infrared touch screen station was mounted (Lafayette Instrument Company, Lafayette, IN). Access to the testing chamber was restricted except for the duration of time during which cognitive testing occurred. During testing, the chamber was accessed via a small doorway, and marmosets could freely enter and exit the chamber. Marmosets earned liquid rewards (e.g., apple juice) that were dispensed into a “sink” below the touchscreen. Potential locations in which stimuli could appear during cognitive testing were indicated by cutouts in a plastic panel that was placed in front of the screen.

#### Software

Animal Behavior Environment Test (ABET) Cognition software (Lafayette Instrument Company, Lafayette, IN) was used to program and control all aspects of the tasks. Measures such as trial number, correct and incorrect responses, the location on the screen of the rewarded choice and the chosen location, and response latency were recorded. Black and white stimuli that were used for cognitive testing consisted of simple shapes and lines displayed on a black background (see Figures 1A and 3A for examples of stimuli).

**Figure 1.**
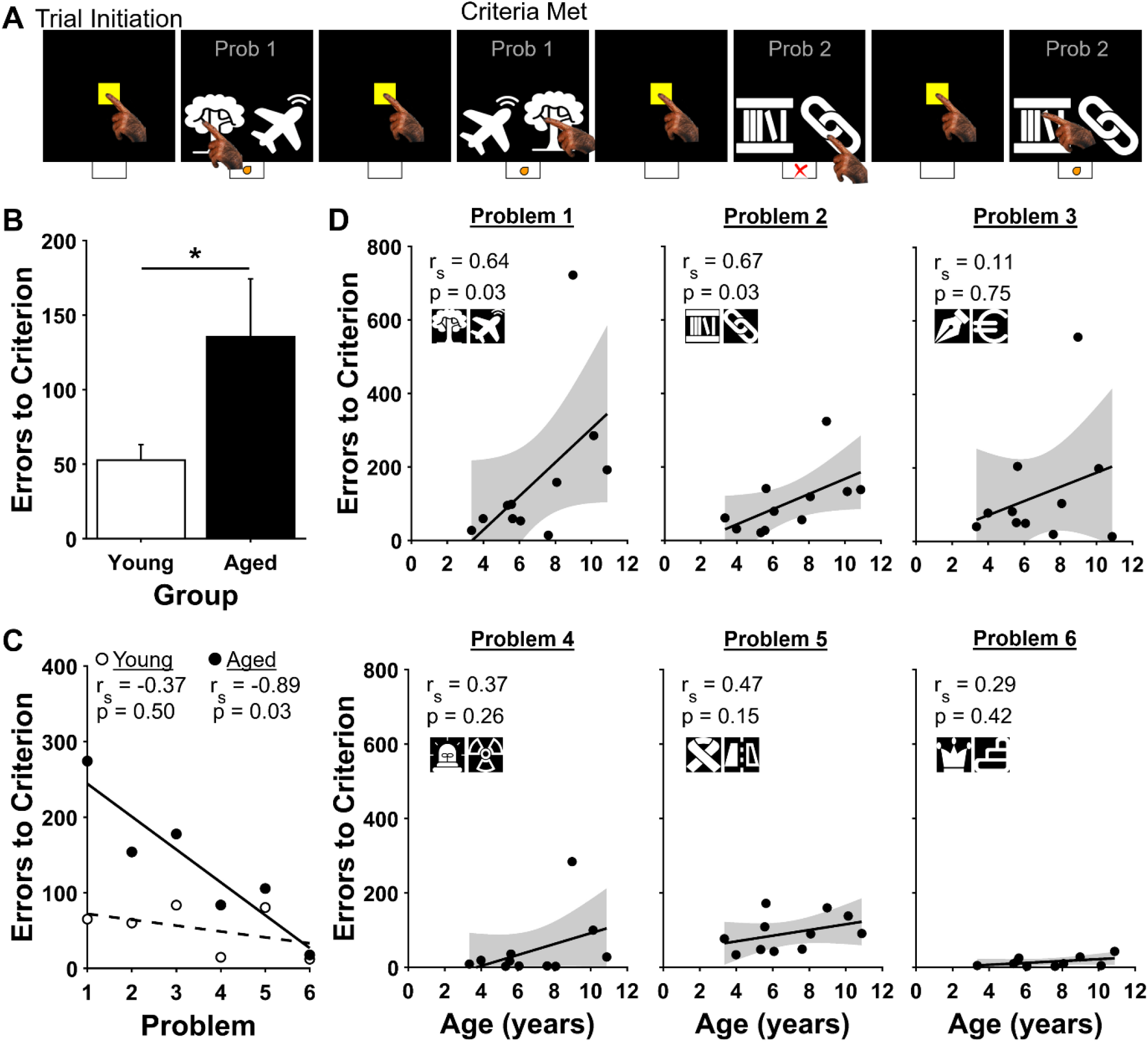
Simple Discrimination (SD) task. (A) Four example SD trials. Below each panel, correct and incorrect choices are indicated by an orange dot and red X, respectively. The first two examples show the last two trials of Problem 1. The next two examples show the first two trials of Problem 2. (B) Errors to criterion by age group. Aged marmosets made more errors to criterion than young marmosets when performance was averaged across all six SD Problems. Bars show mean ± SEM. *p<0.05 (C) Correlations between errors to criterion and Problem by age group. Young marmosets (white circles) performed equally well on all Problems, whereas aged marmosets (black circles) improved their performance over the six SD Problems. (D) Correlations between errors to criterion and age for each of the SD Problems. Strong and significant positive correlations between age and errors to criterion were observed for Problems 1 and 2, but not for Problems 3-6.

### Cognitive Testing

Each of the marmosets in this study had undergone previous cognitive testing, and were therefore proficient at operating the touch screen testing system (Glavis-Bloom et al., 2022).

#### Simple Discrimination (SD)

A Simple Discrimination (SD) task was used to measure marmosets’ capacity to acquire stimulus-reward associations. Each trial of the SD task began when the marmoset touched a yellow square trial initiation stimulus in the center of the screen. Then, two visually distinct black and white stimuli were displayed side-by-side, one randomly assigned to appear on the left, the other on the right. One of the stimuli was predetermined to be the rewarded choice, and the other stimulus was the unrewarded choice. If the marmoset selected the rewarded stimulus, they earned a small liquid reward followed by a 5 second inter trial interval. If the marmoset selected the unrewarded stimulus, or made no selection within 12 seconds (i.e., omitted), no reward was dispensed and a five second time-out period preceded a 5 second inter trial interval. The same pair of stimuli was presented on each trial, with reward contingencies fixed, until the animal demonstrated proficiency by reaching a 90% performance criterion (choosing the rewarded stimulus on 18 of 20 consecutive trials). Once this criterion was achieved for the first SD Problem, the marmosets completed an additional five Problems, one at a time, for a total of six SD Problems. In each of 3-5 testing sessions per week, marmosets performed as many trials as they wanted until the testing session was terminated after 3 hours, or after the marmoset had earned 20 ml of reward, whichever came first. The primary dependent measure of the marmosets’ capacity to acquire stimulus-reward associations was the number of errors to reach the performance criterion on each of the six SD Problems.

#### Serial Reversal (SR)

A Serial Reversal (SR) task was used to measure cognitive flexibility. First, the marmoset completed a single SD Problem to the 90% performance criterion, as described above. Then, without any cue to the marmoset, the reward contingencies were reversed so that the previously rewarded stimulus was now unrewarded and the previously unrewarded stimulus was now rewarded (“Reversal 1”). Marmosets then learned, through trial and error, the new reward contingencies. Once the 90% performance criterion was met on Reversal 1, the reward contingencies were reversed again, reinstating the original reward contingencies (“Reversal 2”). Each time the animal reached criterion, the contingencies were again surreptitiously reversed until the animal completed a total of five Reversals. All other details were identical to those described for the SD task, above. Our measure of cognitive flexibility was the number of errors to criterion on each of the five Reversals, with a smaller number of errors to criterion indicating greater cognitive flexibility.

### Statistical Analyses

Data were analyzed using MATLAB (MathWorks version 2022b). Since the data were not normally distributed, according to a Kolmogorov–Smirnov test, nonparametric statistical tests were used throughout.

Correlations were assessed using Spearman’s rank-order correlations, and effects of age group were analyzed using Mann-Whitney U tests for independent samples. For all statistical tests, p < 0.05 was considered significant.

## RESULTS

### Simple Discrimination (SD)

To assess whether marmosets demonstrate stimulus-reward association deficits with advancing age, we implemented a touch screen version of the Simple Discrimination task in a cohort of animals varying in age from young adult to aged (Figure 1A). Marmosets initiated each trial by touching a yellow square at the center of the screen. This triggered the presentation of two stimuli, one of which was rewarded, the other unrewarded. Marmosets completed six SD Problems, defined as learning to reliably touch the rewarded stimulus when presented with a novel pairing of rewarded and unrewarded stimuli. Training on each new SD Problem proceeded until the monkey achieved proficiency for that pair (90% accuracy calculated over 20 consecutive trials). Trials that were initiated, but where neither stimulus was selected within 12 seconds were excluded from analysis. Each of the marmosets successfully reached criterion on each of the six SD Problems. When performance was averaged across all six Problems, aged marmosets (7 years of age or older) made more errors to criterion than young marmosets (less than 7 years of age), indicating poorer performance (Figure 1B; Mann-Whitney U = 374, p = 0.048). Further, while aged marmosets improved significantly over additional Problems, young animals performed equally well on all Problems, and showed no significant change in performance across them (Figure 1C; aged: r_s_(4) = -0.89, p = 0.03; young: r_s_(4) = -0.37, p = 0.50).

While aged marmosets did learn the SD task, they exhibited more errors to criterion than did young animals. It is an open question whether this reflects an impairment that is specific to learning the new stimulus-reward relationship in each SD problem, or instead reflects an impaired ability to learn the SD task itself. The first of these possibilities predicts impaired learning of each new SD Problem. The latter possibility predicts impairment on early Problems, when the monkey was first learning the SD task, and no impairment on later Problems after the monkey learned the rules of the SD task itself. To distinguish between these two possibilities, we analyzed errors to criterion for each marmoset on each Problem separately. There was a strong and significant positive correlation between age and errors to criterion for the first two Problems, but not for the remaining four (Figure 1D; Problem 1: r_s_(9) = 0.64, p = 0.03; Problem 2: r_s_(9) = 0.67, p = 0.03; Problem 3: r_s_(9) = 0.11, p = 0.75; Problem 4: r_s_(9) = 0.37, p = 0.26; Problem 5: r_s_(9) = 0.47, p = 0.15; Problem 6: r_s_(9) = 0.29, p = 0.42). Thus, while aged animals were impaired on early Problems, they learned the subsequent Problems as efficiently as did young animals.

There are at least two distinct elements the monkeys need to learn in order to perform the SD task accurately. The first of these is to learn the overall rule that governs the structure of the SD task (i.e., “SD task rule”; one stimulus is always rewarded and the other is never rewarded). Second, the monkeys need to apply this rule to the specific stimuli that are unique to each of the SD Problems. That is, for each Problem, the monkey needs to use the SD task rule to successfully acquire a stimulus-reward association. It is an open question whether aged marmosets perform more poorly than young marmosets on the first two SD Problems because of difficulty learning one, the other, or both of these elements.

We defined two phases of the learning curve: an early “Latent Phase” during which the animal performed no better than chance, followed by a “Criterion Phase” during which the animal’s performance was above chance and tended to improve, until the animal reached criterion (Figure 2A). We posit that in the Latent Phase, the monkeys learned the SD task rule, and that the Criterion Phase reflects the period during which the monkeys learn the particular stimulus-reward association that is unique to each Problem. To identify these Phases, we obtained learning curves for each animal and smoothed them with a Robust Locally Weighted Scatterplot Smoothing algorithm over a rolling block of ten trials (Rehse et al., 2016; del Pozo Cruz and del Pozo-Cruz, 2021). Kolmogorov-Smirnov Goodness of Fit Tests were then used to identify the point on each learning curve where performance was significantly above chance (50%) and remained above chance for the rest of the Problem. Trials prior to and including this point are included in the Latent Phase, and trials after this point are included in the Criterion Phase.

**Figure 2.**
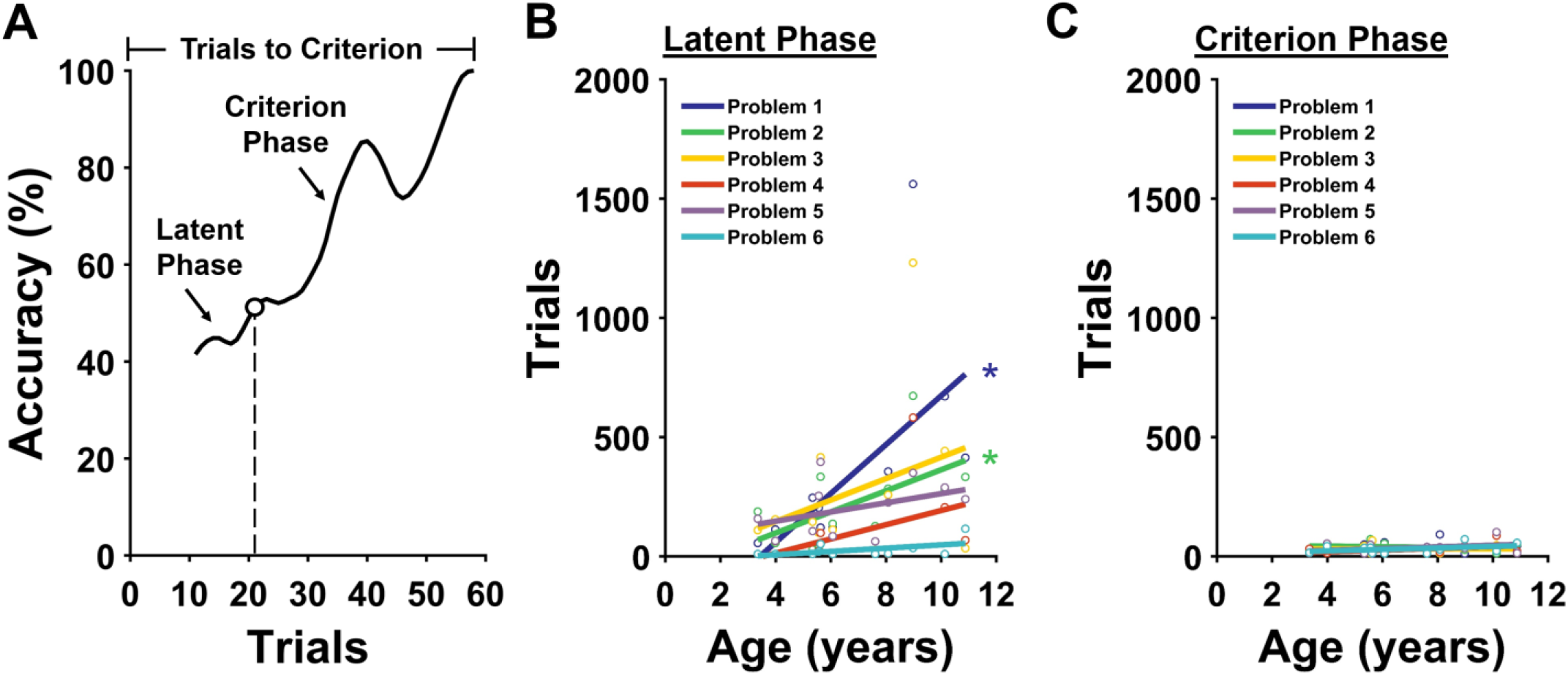
Age-related impaired Simple Discrimination (SD) performance is accounted for by increased duration of the Latent Phase. (A) Representative marmoset learning curve showing performance on one SD Problem. Open circle on the curve marks the point at which performance was significantly above chance (50%) and remained so for the duration of the Problem. (B-C) Correlations between age and number of trials in the (B) Latent Phase and (C) Criterion Phase. These panels show that the duration of the Latent Phase accounts for the age-related impairment on Problems 1 and 2. *p<0.05

**Figure 3.**
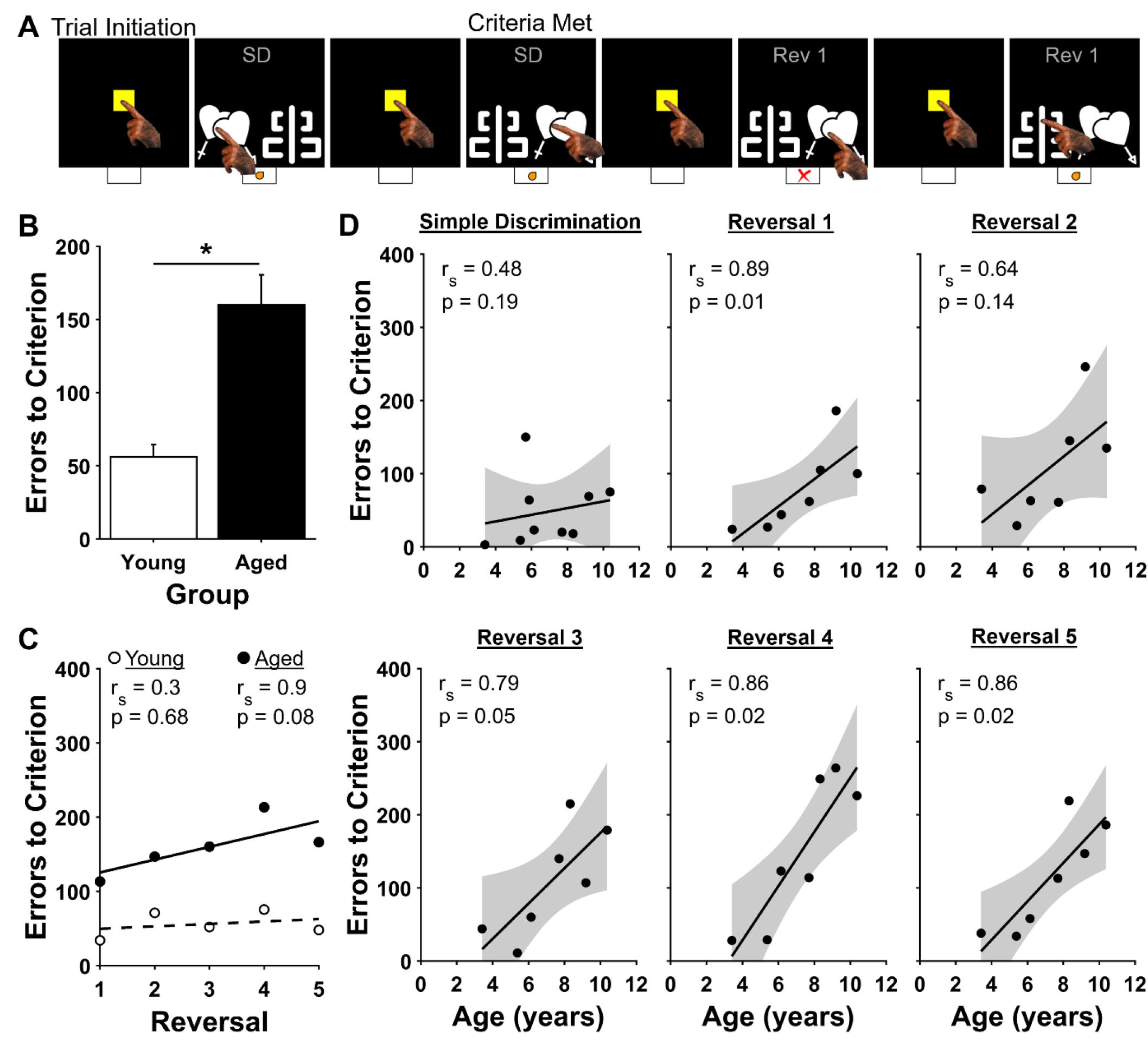
Serial Reversal (SR) task. (A) Example of four SR task trials. Below each panel, correct and incorrect choices are indicated by an orange dot and red X, respectively. The first two examples show the last two trials of the SD Problem that precedes the first Reversal. The next two examples show the first two trials of Reversal 1. (B) Errors to criterion by age group. Aged marmosets made more errors to criterion than young marmosets when performance was averaged across all five Reversals. Bars show mean ± SEM. *p<0.05 (C) Correlations between errors to criterion and Reversal by age group. Young marmosets (white circles) performed equally well on all Problems, whereas aged marmoset (black circles) performance worsened over the five Reversals. (D) Correlations between errors to criterion and age for the SD Problem and each of the Reversals. Strong positive correlations between advancing age and poorer performance existed for all of the Reversals.

Using this approach, we found that the length of the Criterion Phase did not change as a function of marmoset age for any of the SD Problems (Figure 2C; Table 1). This demonstrates that the marmosets’ ability to learn stimulus-reward associations is conserved with age. On the other hand, the length of the Latent Phase did change as a function of age for the first two SD Problems, but did not for any of the later four (Figure 2B; Table 1). This likely reflects that, as monkeys begin to achieve understanding of the SD task rule, they can apply it to the stimuli in Problem 1. That is, learning of the SD task rule occurs over time, and need not be perfect for the monkey to reach criterion. Therefore, even after achieving criterion on Problem 1, they can, in principle, still improve their understanding of the task rule. Thus, we conclude that aged marmosets perform more poorly than young marmosets on the first two SD Problems because they are impaired in learning the overall rule that governs the structure of the SD task, and that aged marmosets are not impaired in learning stimulus-reward associations.

**Table 1.** Simple Discrimination (SD) statistics for each Problem for each Phase. The length of the Latent Phase changed as a function of age for the first two SD Problems, but not for any of the later four. The length of the Criterion Phase did not change as a function of age for any of the SD Problems.

| Problem | Latent Phase | Criterion Phase |
| --- | --- | --- |
| Problem 1 | $r_s(9) = 0.68, p = 0.02$ | $r_s(9) = -0.21, p = 0.54$ |
| Problem 2 | $r_s(9) = 0.61, p = 0.05$ | $r_s(9) = -0.16, p = 0.63$ |
| Problem 3 | $r_s(9) = 0.17, p = 0.61$ | $r_s(9) = 0.03, p = 0.94$ |
| Problem 4 | $r_s(9) = 0.47, p = 0.15$ | $r_s(9) = 0.30, p = 0.38$ |
| Problem 5 | $r_s(9) = 0.38, p = 0.25$ | $r_s(9) = 0.15, p = 0.66$ |
| Problem 6 | $r_s(9) = 0.45, p = 0.16$ | $r_s(9) = 0.19, p = 0.58$ |

### Serial Reversal (SR)

To assess whether marmosets demonstrate age-related impairment of cognitive flexibility, we administered a Serial Reversal task to a subset of the same animals previously tested on the SD task (Figure 3A). First, marmosets completed one additional SD Problem. Consistent with performance on the last several Problems of the SD task, there was no significant correlation between age and errors to criterion (Figure 3D; r_s_(5) = 0.48, p = 0.19). Next, without any signal to the marmoset, the reward contingencies for the stimuli were reversed such that the previously rewarded stimulus now was unrewarded. Each marmoset completed five of these Reversals, with each of the Reversals using the same stimuli as the SD Problem that was presented at the beginning of SR testing. We used the number of errors committed by subjects on Reversals as a measure of cognitive flexibility, with fewer errors indicating better cognitive flexibility and more errors indicating worse cognitive flexibility (Zeamer et al., 2011a; Weiss et al., 2019). When performance was averaged across all five Reversals, aged marmosets (7 years of age or older) made substantially more errors to criterion than did young marmosets (less than 7 years of age) (Figure 3B; Mann-Whitney U = 11, p < 0.00001). Further, whereas young marmosets’ performance was stable across Reversals, aged marmosets’ performance worsened across the Reversals, as revealed by a strong positive correlation between Reversal number and errors to criterion that approached significance (Figure 3C; Young: r_s_(3) = 0.03, p = 0.68; Aged: r_s_(3) = 0.90, p = 0.08).

As we did for the SD task, we next analyzed errors to criterion for each marmoset on each of the Reversals separately. There were strong positive correlations between age and errors to criterion that persisted for all of the Reversals (Figure 3D; Reversal 1: r_s_(5) = 0.89, p = 0.01; Reversal 2: r_s_(5) = 0.64, p = 0.14; Reversal 3: r_s_(5) = 0.79, p = 0.05; Reversal 4: r_s_(5) = 0.86, p = 0.02; Reversal 5: r_s_(5) = 0.86, p = 0.02). Together, these results demonstrate that cognitive flexibility worsens with increasing age, and does not improve with experience.

To successfully perform Reversals, marmosets need to switch their response strategies to avoid the previously rewarded stimulus and choose the previously unrewarded stimulus (Weiss et al., 2019). There are at least three elements to this process. First, the animal needs to learn that a previously established behavioral response is no longer satisfactory. That is, that the previously rewarded stimulus, when selected, no longer yields a reward. Second, the animal needs to suppress a behavioral response to the previously rewarded stimulus, thereby resulting in some selection of the newly rewarded stimulus. Third, the subject must successfully acquire the new stimulus-reward association. It is an open question which of these elements contributes to impaired performance of the aged animals.

We defined three distinct Phases of the marmosets’ learning curves for each Reversal: a “Transition Phase” during which marmosets performed at or below 10% accuracy, followed by a “Dynamic Phase” during which performance gradually increased to above chance, and a “Criterion Phase” during which the animal’s performance was above chance and tended to improve, until the animal reached criterion (Figure 4A). We posit that in the Transition Phase, monkeys learn that their previously established behavioral response no longer yields rewards, and that the Dynamic Phase reflects the period during which the monkey begins to suppress selection of the previously rewarded stimulus and select the newly rewarded stimulus at least half of the time. Finally, we posit that during the Criterion Phase, monkeys learn the stimulus-reward association for the Reversal. As with the SD task, to identify these Phases, learning curves from the Reversals were smoothed with a Robust Locally Weighted Scatterplot Smoothing algorithm over a moving block of ten trials. Then, Kolmogorov-Smirnov Goodness of Fit Tests were used to identify the point at which performance was significantly greater than 10% and remained above 10% for the remainder of the Reversal (i.e., the boundary between the Transition and Dynamic Phases), and the point at which performance was significantly greater than chance (50%) and remained above chance for the remainder of the Reversal (i.e., the boundary between the Dynamic and Criterion Phases).

**Figure 4.**
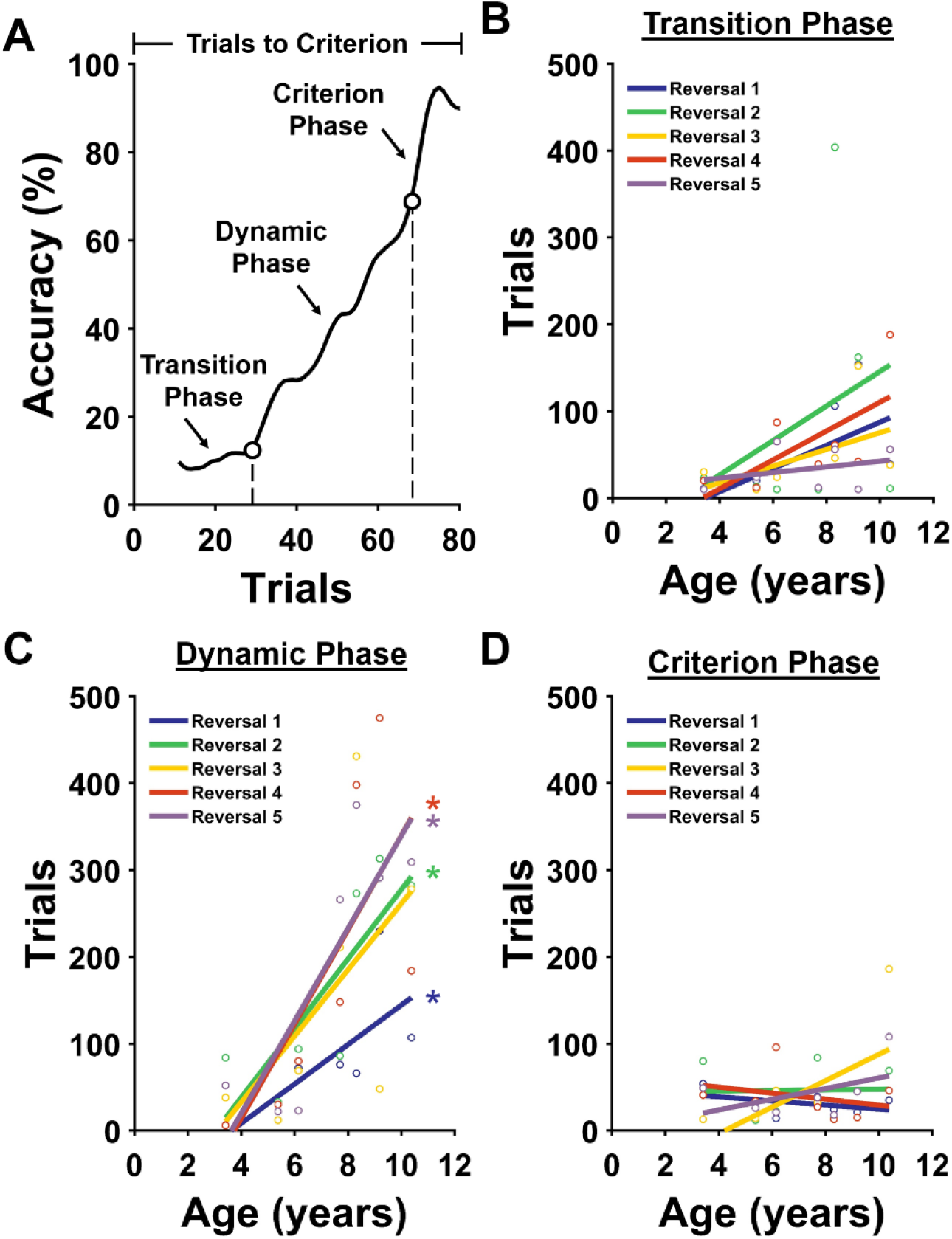
Age-related impaired Serial Reversal (SR) performance is accounted for by increased duration of the Dynamic Phase. (A) Representative marmoset learning curve showing performance on one Reversal. The first trial on the x-axis corresponds to the first trial after the reward contingencies were switched. The lower open circle on the curve marks the point at which performance was statistically above 10% accuracy and remained so for the duration of the Reversal. Trials prior to this point are included in the Transition Phase. The higher open circle indicates the point at which performance exceeded chance levels (50%) and remained above chance for the remainder of the Reversal. Trials above this point are included in the Criterion Phase. Trials between the end of the Transition Phase and beginning of the Criterion Phase are included in the Dynamic Phase. (B-D) Correlations between age and number of trials in the (B) Transition Phase, (C) Dynamic Phase, and (D) Criterion Phase. These panels show that the duration of the Dynamic Phase accounts for the overall age-related impairment in cognitive flexibility. *p<0.05

Using this approach, we found that the length of the Dynamic Phase increased significantly as a function of age (Figure 4C; Table 2), whereas the lengths of the Transition and Criterion Phases did not vary significantly as a function of age (Figures 4B&D; Table 2). Taken together, these results demonstrate that aged marmosets have impaired cognitive flexibility that can be attributed to impaired suppression of the previously rewarded behavioral response. Further, as was found on the SD task, once marmosets reliably perform above chance levels, there is no systematic change with age.

**Table 2.** Serial Reversal (SR) statistics for each Reversal for each Phase. The length of the Dynamic Phase significantly increased as a function of age, whereas the lengths of the Transition and Criterion Phases did not vary significantly as a function of age.

| Reversal | Transition Phase | Dynamic Phase | Criterion Phase |
| --- | --- | --- | --- |
| Reversal 1 | $r_s(5) = 0.67, p = 0.12$ | $r_s(5) = 0.86, p = 0.02$ | $r_s(5) = -0.29, p = 0.56$ |
| Reversal 2 | $r_s(5) = 0.13, p = 0.79$ | $r_s(5) = 0.89, p = 0.01$ | $r_s(5) = 0.07, p = 0.91$ |
| Reversal 3 | $r_s(5) = 0.68, p = 0.11$ | $r_s(5) = 0.68, p = 0.11$ | $r_s(5) = 0.50, p = 0.25$ |
| Reversal 4 | $r_s(5) = 0.71, p = 0.09$ | $r_s(5) = 0.89, p = 0.01$ | $r_s(5) = -0.25, p = 0.59$ |
| Reversal 5 | $r_s(5) = 0.02, p = 0.67$ | $r_s(5) = 0.79, p = 0.04$ | $r_s(5) = 0.21, p = 0.66$ |

### Control Analyses

The animals studied here have previously been found to exhibit age-related changes in motivation and motor speed (Glavis-Bloom et al., 2022). We used trial completion rate to assess age-related changes in motivation on the SD and SR tasks. Each of the marmosets completed more than 95% of the trials they initiated on each task, and trial completion rate did not significantly change as a function of age (SD: r_s_(9) = 0.43, p = 0.19; SR: r_s_(5) = 0.18, p = 0.71). Thus, the age-related changes reported above cannot be understood as stemming from age-related changes in motivation. Next, we used correct choice response latency to assess age-related motor slowing in the context of the SD and SR tasks. We found no significant associations between age and correct choice response latency during the blocks of 20 trials on which criterion were met on either task (SD: r_s_(9) = -0.05, p = 0.88; SR: r_s_(5) = -0.21, p = 0.66). This demonstrates that the age-related impairments shown above do not reflect age-related changes in motor speed.

## DISCUSSION

In this study, we characterized stimulus-reward association learning and cognitive flexibility in young adults to geriatric marmosets using a Simple Discrimination (SD) and a Serial Reversal (SR) task, respectively. In the SD task, older marmosets were impaired at learning the SD task rule relative to younger animals. Once this rule was learned, however, aged marmosets were unimpaired in their ability to learn stimulus-reward associations. On the SR task, older marmosets showed remarkably impaired cognitive flexibility relative to younger animals. Accumulated proactive interference from serial reversals exacerbated this age-related impairment.

### Comparison with prior simple discrimination tasks in non-human primates

Our work aligns well with previously published studies that utilized SD tasks in aging non-human primates. First, consistent with prior studies we find that the ability to form stimulus-reward associations remains intact in aging marmosets compared to young (Bartus et al., 1979; Bachevalier et al., 1991; Lai et al., 1995; Burke et al., 2011; Munger et al., 2017; Sadoun et al., 2019; Gray et al., 2023). Second, some prior studies, and our work reported here, find that any age-related impairment detected by the SD task is accounted for by performance on early trials, when animals are first learning the rule that governs the structure of the SD task. This learning impairment manifests in aged animals in the form of a prolonged period of time in which animals’ performance is at levels approximating chance (Voytko, 1999; Zeamer et al., 2011a; Munger et al., 2017; Gray et al., 2023).

Our work extends these findings by revealing that these impairments are transient, disappearing completely once the aged animals have learned the structure of the task. Specifically, we find that, in older marmosets, the duration of chance-like performance decreases with SD task experience. We also find that once marmosets reliably perform above the levels expected by chance, the rate at which they reach criterion-levels of performance does not change as a function of age. Together, these results demonstrate that the age-related SD task impairment is attributable to impairment in “learning-to-learn” (Harlow, 1949), not an impairment in the learning of particular stimulus-reward associations. Our behavioral findings align with what we would expect from previous neurophysiological and computational studies that report learning-to-learn critically depends on the prefrontal cortex (Cromer et al., 2011; Goudar et al., 2023), and the fact that the prefrontal cortex undergoes morphological and functional changes early in the aging process (Upright and Baxter, 2021).

Our conclusions do differ from one previous study that reported impaired stimulus-reward association learning in aged marmosets (Rothwell et al., 2022). Several differences between this study and ours may account for the difference in findings. Rothwell and colleagues tested a group of adult marmosets on three SD Problems, once per year, for four years. During the first three years of the study, marmosets did not show an impairment (Workman et al., 2019). However, when tested in year four, an impairment was observed across all animals. This impaired performance may be attributable to a change in experimental design that coincided with the drop in performance. Whereas the stimuli used in years one, two, and three were visually distinct in multiple dimensions, changing both in color and in shape, stimuli used in year four were more similar to one-another, varying only in the shape dimension, while matching in color. Since stimulus similarity is known to affect discriminability (Zeamer et al., 2011b), we controlled for this by using only black and white stimuli throughout our study. Additionally, while marmosets in the Rothwell et al. study completed three SD Problems each year, the data were analyzed by collapsing performance across each of the Problems completed in a given year. In contrast, we administered six SD Problems to our marmosets, and analyzed performance on each Problem separately. This enabled us to observe how performance changed not only as a function of age, but also as a function of experience. We found that the rate of learning the rule that governs the structure of the SD task changed as a function of age, but that performance across the age range was equivalent given experience. Therefore, we posit that Rothwell et al’s results may be stimulus-driven or due to a learning deficit on SD rather than a stimulus-reward association deficit.

### Comparison with prior reversal learning tasks in non-human primates

Our finding that older marmosets have impaired cognitive flexibility compared to younger marmosets aligns with a large body of existing macaque and marmoset literature (Lai et al., 1995; Voytko, 1999; Gray et al., 2017, 2018; Munger et al., 2017; Sadoun et al., 2019). Our work extends these findings by demonstrating that aged marmosets are particularly susceptible to the effects of proactive interference. By analyzing performance on discrete segments of the learning curves, we show that the impaired cognitive flexibility of older marmosets is driven exclusively by extended durations of the Dynamic Phase, as compared to younger marmosets. This finding aligns with what is expected if aged animals had difficulty disengaging from previously formed stimulus-reward associations, indicating increased susceptibility to proactive interference compared to young (Gray et al., 2017). Overcoming proactive interference is a major function of the prefrontal cortex (Jha et al., 2004; Burke et al., 2009; Cromer et al., 2011), which is known to dysfunction with aging, and therefore offers a neuroanatomical explanation for age-related susceptibility to proactive interference (Morrison and Baxter, 2012; Upright and Baxter, 2021).

To escalate the demands on cognitive flexibility and probe the limits of this domain as a function of age we implemented a serial reversal paradigm (Dickinson and Mackintosh, 1978; Hassett and Hampton, 2017). While there is a large body of non-human primate literature that has utilized serial reversals to assess cognitive flexibility (Roberts et al., 1990; Izquierdo et al., 2004; Rudebeck and Murray, 2011; Jackson et al., 2019), ours is the first to assess the consequences of aging using this paradigm. We found that, whereas performance of young marmosets was stable across Reversals, aged marmoset performance progressively worsened. We posit that this exacerbation of the cognitive flexibility impairment is due to accumulated proactive interference from the serial nature of the Reversals.

## Acknowledgments

This research was supported by an AHA-Allen Initiative in Brain Health and Cognitive Impairment award made jointly through the American Heart Association and The Paul G. Allen Frontiers Group: 19PABH134610000AHA, a National Institutes of Health grant 1R21AG068967-01, grants from the Larry L. Hillblom Foundation and the Don and Lorraine Freeberg Foundation, and the Fiona and Sanjay Jha Chair in Neuroscience. We thank Katie Williams for assistance in the care of the marmosets and technical support.

## Summary

Here we have established that aged marmosets have conserved stimulus-reward association learning, but show transient impairment in learning-to-learn. Furthermore, aged marmosets have impaired cognitive flexibility driven by susceptibility to proactive interference. Accumulated proactive interference significantly exacerbates cognitive flexibility impairment. Each of these impairments are in domains critically supported by the prefrontal cortex. Therefore, our findings support prefrontal cortical dysfunction as a prominent feature of neurocognitive aging and position the marmoset as a key model for determining the neural underpinnings of age-related prefrontal cortical dysfunction.

